# Differences in pulmonary innate lymphoid cells are dependent on mouse age, sex and strain

**DOI:** 10.1101/2020.10.25.354464

**Authors:** Svenja Loering, Guy J. M Cameron, Nirmal P Bhatt, Gabrielle T Belz, Paul S Foster, Philip M Hansbro, Malcolm R Starkey

## Abstract

Innate lymphoid cells (ILC) are resident in the lung and are involved in both the maintenance of homeostasis and the pathogenesis of respiratory diseases. In this study, murine lung ILC were characterised using flow cytometry and the impact of mouse age, sex and strain were assessed. Lung ILC were found as early as postnatal day 4 and numbers peaked at 2 weeks, and then decreased as the lung matured. During postnatal lung development, ILC expressed differential amounts of ILC2-associated cell surface antigens including ST2, CD90.2 and ICOS. Using *Il5*^venus^*Il13*^td-tomato^ dual reporter mice, neonates were found to have increased constitutive IL-13 expression compared to adult mice. Neonates and adults had similar ratios of IL-5^+^CD45^+^ leukocytes, however, these cells were mostly composed of ILC in neonates and T cells in adults. Sex-specific differences in ILC numbers were also observed, with females having greater numbers of lung ILC than males in both neonatal and adult mice. Female lung ILC also expressed higher levels of ICOS and decreased KLRG1. Mouse strain also impacted on lung ILC with BALB/c mice having more ILC in the lung and increased expression of ST2 and ICOS compared with C57BL/6J mice. Collectively, these data show that lung ILC numbers, cell surface antigen expression, IL-5 and IL-13 levels differed between neonatal and adult lung ILC. Additionally, cell surface antigens commonly used for ILC2 quantification, such as ST2, CD90.2, and ICOS, differ depending on age, sex and strain and these are important considerations for consistent universal identification of lung ILC2.

## Introduction

In mice, innate lymphoid cells (ILC) are lineage marker (Lin) and T cell receptor (TCR) negative lymphocytes that can be broadly classified into three groups based on their cytokine and transcription factor expression^1,2^. Group 1 ILC (ILC1) are characterised by their expression of interferon (IFN)-γ and T-bet^1,2^. ILC2 express type 2 cytokines such as interleukin (IL)-5 and IL-13 and the transcription factor GATA3, whilst ILC3 express IL-17 and/or IL-22, in addition to RORγt^1,2^. ILC2 are the predominant ILC subtype in the lung, where they contribute to the maintenance of lung homeostasis and to the pathogenesis of respiratory diseases such as asthma, chronic obstructive pulmonary disease and influenza virus infection^1,3,4^.

Recently, it has been shown that several type 2 immune cells, including ILC2, are present in mouse lungs shortly after birth^5–8^. These cells peak in number during the first two weeks of life and decrease by adulthood^5–8^. The first few weeks of life are associated with the development and maturation of the mouse lung^9^. Unlike other organs, the lung starts to develop *in utero*, and continues after birth^9^. Lung development is separated into five stages; embryonic, pseudoglandular, canalicular, saccular and alveolar^9^. The saccular stage begins *in utero* and continues postnatally, followed by the alveolar stage^9^. Both stages are characterised by maximal remodelling, as at birth, the liquid-filled lung is suddenly exposed to air^9^. The first breath is thought to induce the spontaneous release of IL-33, causing an increase in lung-resident type 2 immune cells^5,6^. These include ILC2, which are activated and proliferate in response to IL-33 *via* the IL-33 receptor ST2^3^.

The increase in ILC2, and, as recently identified, ILC3, during postnatal lung remodelling suggests that ILC may play a role in lung developmental processes^1,5–8,10,11^. In this study, lung ILC were characterised at distinct time points during postnatal lung development, with a focus on neonates at postnatal day (P) 10 and adult mice aged between 7-9 weeks. We show that neonates have higher numbers of lung resident ILC than adults, with increased expression of the ILC2-associated markers ST2, ICOS and CD90.2. Sex-specific differences were also observed with females having more lung ILC than males, with higher expression of ICOS but lower expression of KLRG1. Mouse strain also impacted on lung ILC numbers with BALB/c mice having more ILC than C57BL/6J mice.

## Results

### Neonates have increased numbers of lung ILC and different ILC cell surface antigen expression compared to adult mice

In order to determine the number of lung ILC during postnatal lung development, C57BL/6J mice were sacrificed at several time points between postnatal day (P) 4 and 21. Lung ILC were characterised as CD45^+^ TCR^−^ (TCRβ^−^TCRγδ^−^CD4^−^CD8^−^) Lin^−^ (CD11b^−^B220^−^Ly6a/e^−^ NK1.1^−^CD3^−^Ter-119^−^) CD2^−^IL-7R^+^ cells (Figure 1a). There were clear lung ILC populations present by P4, and ILC numbers peaked at P14, before decreasing by P21 (Figure 1b-g). Cell surface antigen expression of lung ILC2-associated markers varied during the postnatal time points assessed (Figure 1h-l). ST2 expression peaked at P7 (Figure 1h). CD90.2 expression stayed relatively constant throughout postnatal lung development (Figure 1i), whereas SCA-1 expression was highest between P7-10 (Figure 1j). ICOS peaked at P7 and decreased from P10 (Figure 1k), and KLRG1 expression increased by P7 and stayed relatively stable thereafter (Figure 1l).

**Figure 1:**
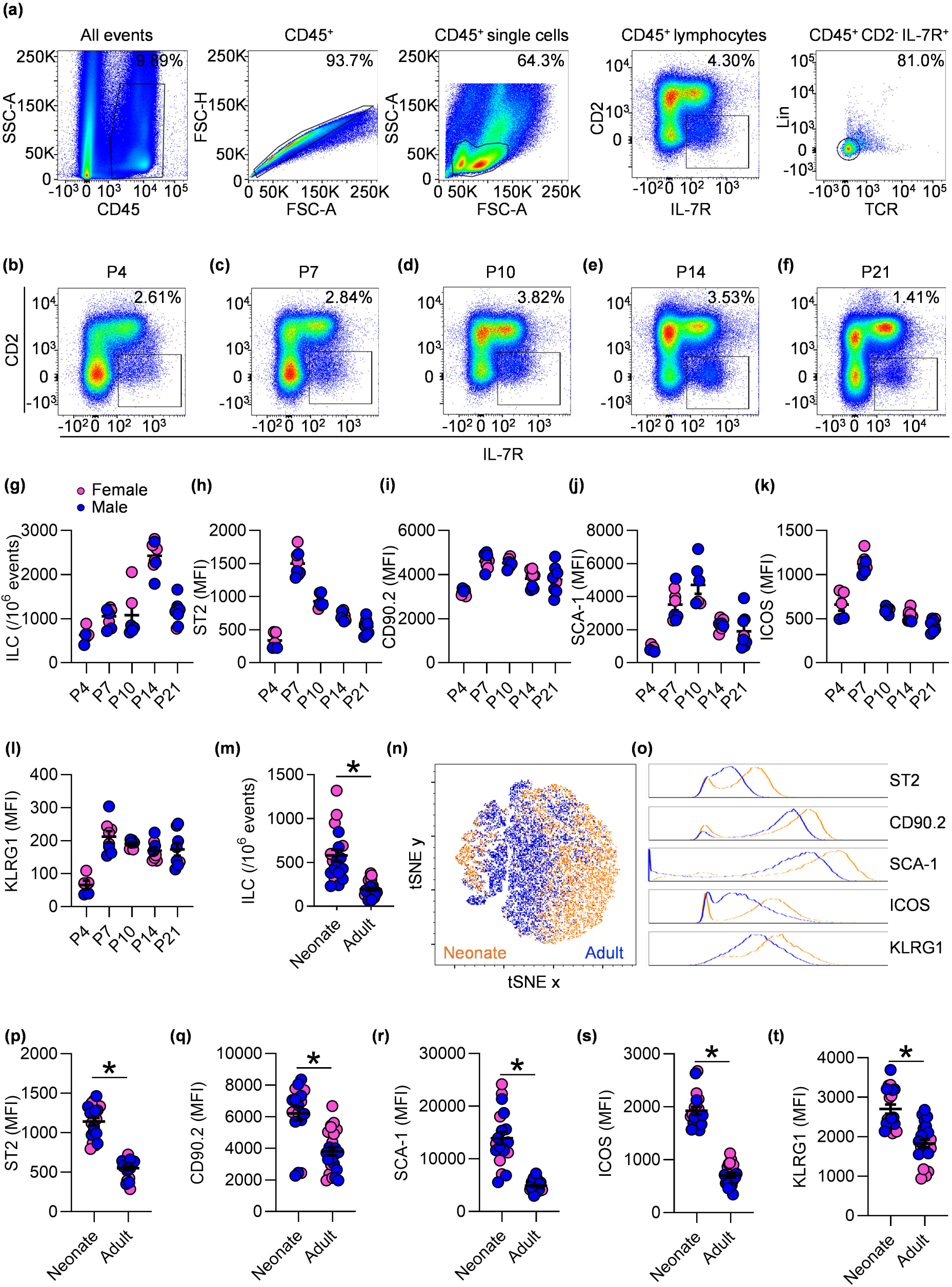
Neonates have increased numbers of lung ILC and different ILC cell surface antigen expression compared to adult mice. **(a)** Representative gating strategy for lung ILC. ILC were characterised as CD45^+^ T cell receptor^−^ (TCR; TCRβ^−^TCRγδ^−^CD4^−^CD8^−^) lineage^−^ (Lin; CD11b^−^B220^−^Ly6a/e^−^NK1.1^−^CD3^−^Ter-119^−^)CD2^−^IL-7R^+^ cells. **(b-f)** Representative pseudocolour plots of CD45^+^ lymphocytes shown as CD2^+^ *versus* IL-7R^+^ from **(b)** postnatal day (P) 4, **(c)** P7, **(d)** P10, **(e)** P14 and **(f)** P21. **(g)** Numbers of lung ILC per 10^6^ events. **(h-l)** Mean fluorescent intensity (MFI) of ILC cell surface marker expression. **(m)** Number of ILC in P10 (neonate) and 7-9-week-old (adult) mice. **(n)** Dimensionality reduction of ILC in relation to ST2, CD90.2, SCA-1, ICOS and KLRG1 using t-Distributed Stochastic Neighbour Embedding (tSNE). tSNE x and tSNE y represent the two new parameters. Orange: neonate at P10, blue: adult (7-9 weeks). (**o**) Histograms of ILC2-associated cell surface antigens. (**p-t**) Quantified MFI from (n). Pink female, blue male mice. Data are presented as the mean ± the standard error of the mean (s.e.m). Data are representative of 2-5 independent experiments. Statistical significance was calculated using non-parametric unpaired Mann-Whitney t test. * P<0.05.

Next, two representative time points were chosen to assess the differences between neonate (P10) and adult (7-9 weeks) lung ILC. There were significantly higher proportions of ILC in neonates compared to adult lungs relative to the total number of events counted (Figure 1m). t-distributed stochastic neighbour embedding (tSNE) is an unbiased algorithm used to cluster populations based on the expression of cell surface markers. Neonatal and adult ILC clustered in distinct populations when analysing the cells for the expression of ST2, CD90.2, SCA-1, ICOS and KLRG1 (Figure 1n). Neonatal ILC showed higher expression of all cell surface antigens compared to adult ILC (Figure 1o-t).

### Female mice have increased numbers of lung ILC compared to age-matched males

Our initial data indicated sex-specific differences in the number of lung resident ILC, with female mice having increased ILC compared to male mice (Figure 1m). Splitting the data based on sex showed that female mice had increased ILC numbers compared to male mice in both neonates and adults (Figure 2a-e). Clustering analysis of cell surface antigen expression did not show sex-specific differences in neonates (Figure 2f-g). Adult mice showed less overlap between the sexes, with increased expression of CD90.2 and ICOS, as well as decreased expression of KLRG1 in females compared to males (Figure 2h-i). Quantification of the mean fluorescent intensity (MFI) of the cell surface receptors on lung ILC revealed no differences between the sexes in terms of ST2, CD90.2, and SCA-1 expression in either age group (Figure 2j-l). ICOS MFI was increased in females of both ages compared to age-matched male mice (Figure 2m). KLRG1 was increased in adult males compared to adult females, whereas in neonates KLRG1 expression was comparable between the sexes (Figure 2n).

**Figure 2:**
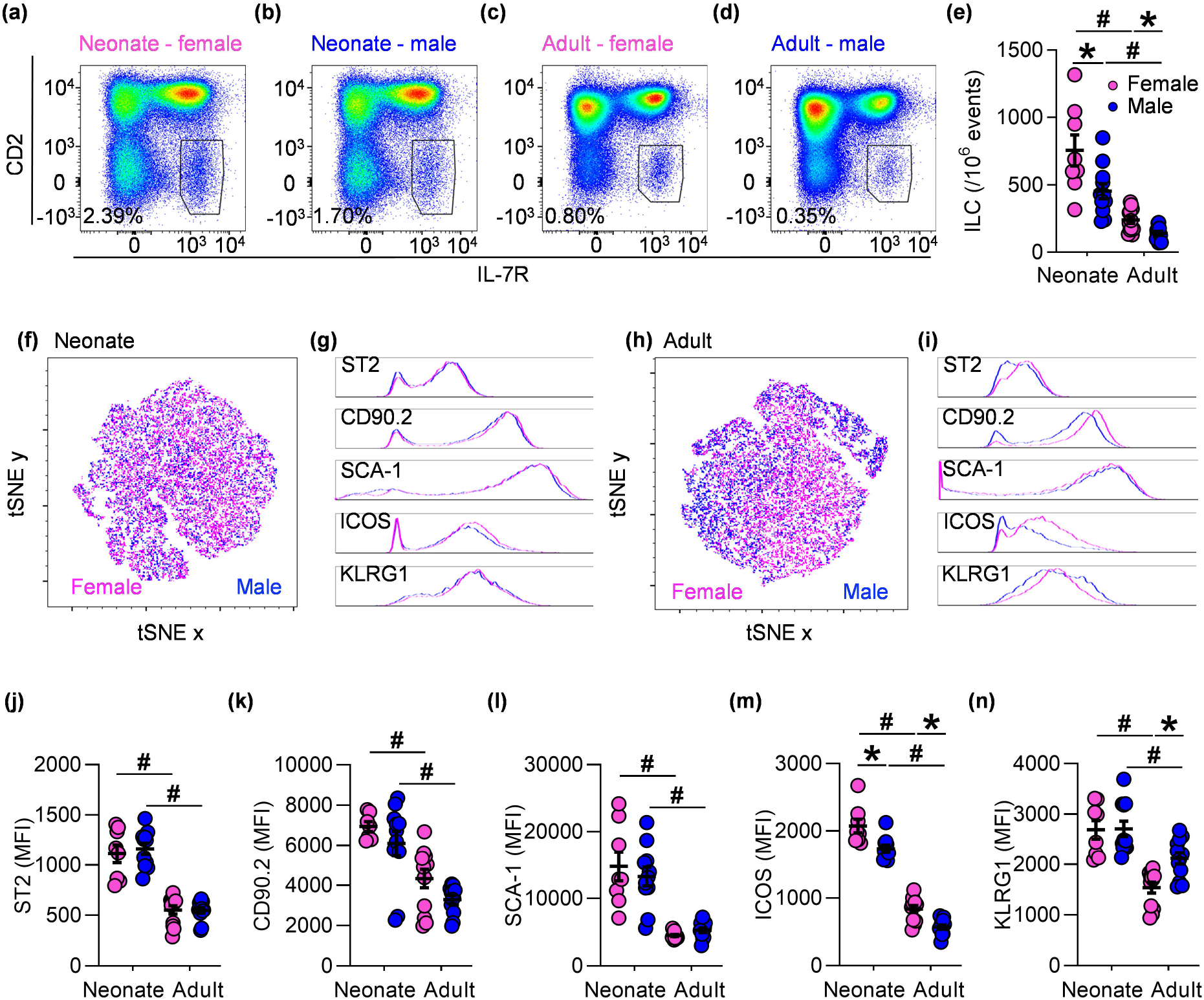
Female mice have increased numbers of lung ILC compared to age-matched males. **(a-d)** Representative pseudocolour plots of CD45^+^ lymphocytes shown as CD2^+^ *versus* IL-7R^+^ for **(a)** female neonate, **(b)** male neonate, **(c)** female adult, and **(d)** male adult mice. **(e)** Quantification of ILC (CD45^+^ T cell receptor^−^ (TCR; TCRβ^−^TCRγδ^−^CD4^−^CD8^−^) lineage^−^ (Lin; CD11b^−^B220^−^Ly6a/e^−^NK1.1^−^CD3^−^Ter-119^−^) CD2^−^IL-7R^+^ cells) in mice from postnatal day (P) 10 and adult mice (7-9 weeks). **(f)** Dimensionality reduction of neonatal ILC relating to ST2, CD90.2, SCA-1, ICOS and KLRG1 using t-Distributed Stochastic Neighbour Embedding (tSNE). tSNE x and tSNE y represent the two new parameters. **(g)** Histograms of ILC2-associated cell surface antigens. **(h)** tSNE and **(i)** corresponding histograms in adult ILC. **(j-n)** Quantified mean fluorescent intensities (MFI). Pink female, blue male mice. Data are presented as the mean ± the standard error of the mean (s.e.m). Data are representative of 2-4 independent experiments. Statistical significance was calculated using non-parametric unpaired Mann-Whitney t test. P<0.05 compared to neonate (*) or female (#).

### Differential expression of the type 2 cytokines IL-5 and IL-13 in neonatal compared to adult lung ILC

ILC2 are known to produce the type 2 cytokines IL-5 and IL-13^5,7,12,13^. To investigate the expression of these cytokines in the neonatal and adult lung, CD45^+^ lymphocytes were analysed for their expression of IL-5 and IL-13 using *Il5*^venus/+^*Il13*^td-tomato/+^ dual reporter mice^14,15^. In neonates, the majority of CD45^+^ lymphocytes were double negative for both cytokines (97.9%), a proportion were double-positive for both IL-5 and IL-13 (0.99%), and minor fractions expressed IL-5 (0.29%) or IL-13 (0.81%; Figure 3a-b). Adults had a higher total number of CD45^+^ cells, with greater IL-5 expression (0.28%) than IL-13 expression (0.19%) and very few double positive cells (0.029%; Figure 3c-d). There were similar total numbers of IL-5^+^CD45^+^ cells at both ages (Figure 3e-g), but neonates had higher IL-5 MFI compared to adults (Figure 3h). IL-13 expression was greater in neonates than adults (Figure 3i-l). Further analysis of the IL-5^+^CD45^+^ cells for their expression of Lin markers and TCR showed that in neonates, most IL-5^+^CD45^+^ cells were Lin^−^TCR^−^ ILC (68.7%) (Figure 3m-n). In adult mice, Lin^+^ and/or TCR^+^ cells made up major proportions of IL-5^+^CD45^+^ cells (Figure 3o-p). In neonates, the majority of IL-13^+^CD45^+^ cells were Lin^−^TCR^−^ ILC (77.6%) (Figure 3q-r), while in adults IL-13^+^CD45^+^ cells comprised both Lin^−^TCR^−^ (57.7%) and Lin^−^ TCR^+^ (34.3%; Figure 3s-t).

**Figure 3:**
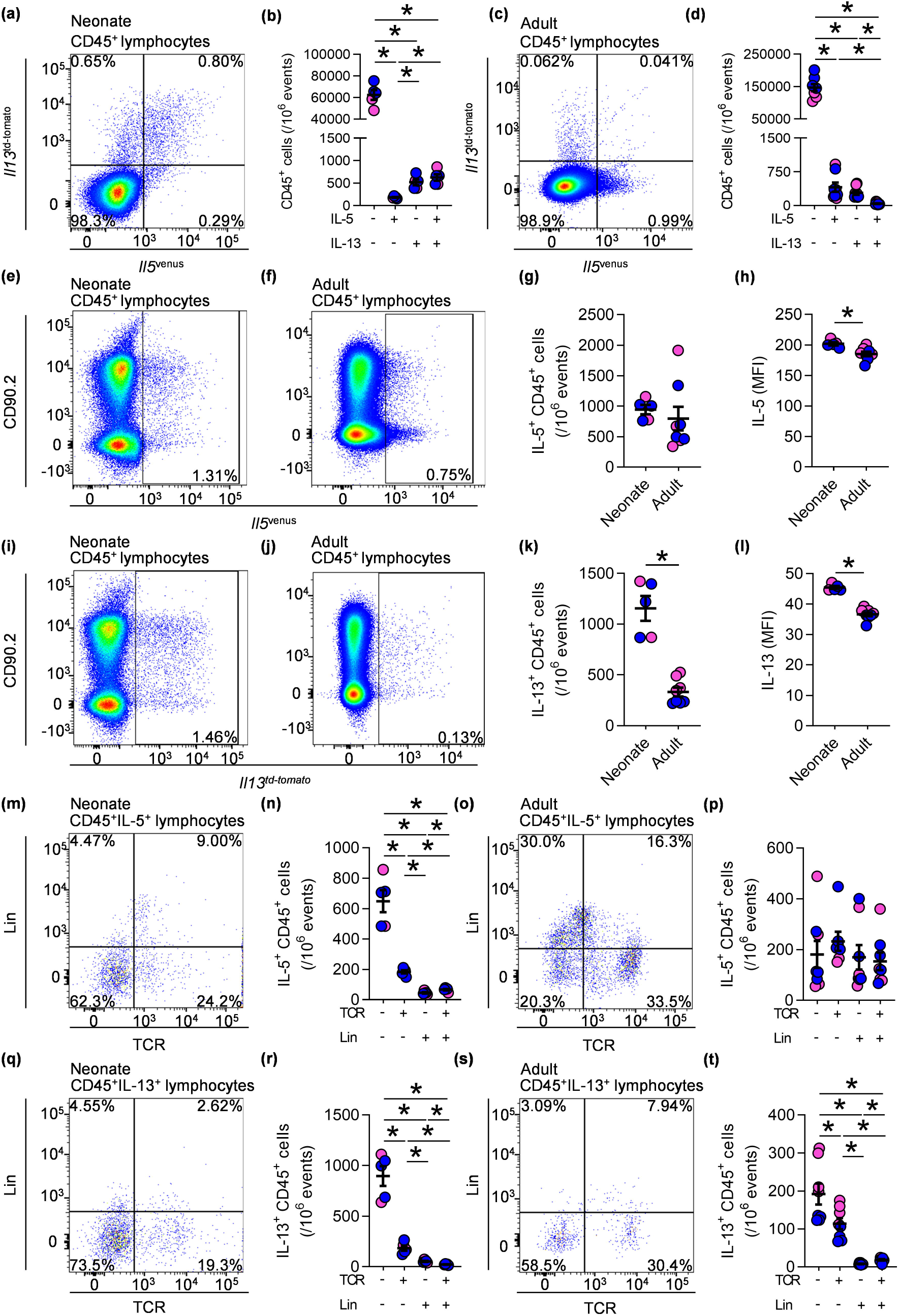
Differential expression of the type 2 cytokines IL-5 and IL-13 in neonatal lung ILC compared to adult lung ILC. **(a)** Representative pseudocolour plot of CD45^+^ lymphocytes shown as *Il5*^venus^ *versus Il13*^td-tomato^ in *Il5*^venus/+^*Il13*^td-tomato/+^ dual reporter mice in neonatal lungs (postnatal day (P) 10). **(b)** Quantification of **a**. **(c)** Representative pseudocolour plot of CD45^+^ lymphocytes shown as *Il5*^venus^ *versus Il13*^td-tomato^ and **(d)** quantification of CD45^+^ lymphocytes in adult lungs. **(e-f)** Representative pseudocolour plots of IL-5^+^ CD45^+^ lymphocytes shown as *Il5*^venus^ *versus* CD90.2 in **(e)** neonates and **(f)** adults. **(g)** Quantification of IL-5^+^ CD45^+^ lymphocytes. **(h)** Mean fluorescent intensity (MFI) of IL-5 in CD45^+^ lymphocytes. **(i-j)** Representative pseudocolour plots of IL-13^+^ CD45^+^ shown as *Il13*^td-tomato^ *versus* CD90.2 in **(i)** neonatal and **(j)** adult mouse lungs. **(k)** Quantification of IL-13^+^ CD45^+^ lymphocytes. **(l)** MFI of IL-13 in CD45^+^ lymphocytes. **(m)** Representative pseudocolour plot of IL-5^+^ CD45^+^ lymphocytes shown as T cell receptor (TCR; TCRβ^−^γδ^−^CD4^−^CD8^−^) *versus* lineage (Lin; CD11b^−^Ly6a/e^−^B220^−^CD3^−^Ter-119^−^NK1.1^−^) markers and **(n)** quantification of IL-5^+^ CD45^+^ lymphocytes in neonates. **(o)** Representative pseudocolour plot and **(p)** quantification of IL-5^+^ CD45^+^ lymphocytes in adults. **(q)** Representative pseudocolour plot of IL-13^+^ CD45^+^ lymphocytes shown as TCR vs Lin and **(r)** quantification of IL-13^+^ CD45^+^ lymphocytes in neonates. **(s)** Representative pseudocolour plot and **(t)** quantification of IL-13^+^ CD45^+^ lymphocytes in adult lungs. Pink female, blue male mice. Data are presented as the mean ± the standard error of the mean (s.e.m). Data are representative of 2-4 independent experiments. Statistical significance was calculated using unpaired and non-parametric Mann-Whitney t test. * P<0.05.

### BALB/c mice have increased numbers of lung ILC and differential expression of ILC2-associated cell surface markers compared to C57BL/6J mice

Analysis of lung ILC numbers in *Il5*^venus/+^*Il13*^td-tomato/+^ BALB/c mice indicated that they may have increased numbers of ILC compared to C57BL/6J mice (data not shown). To confirm this difference, lung ILC were quantified in wild type BALB/c and C57BL/6J mice. BALB/c mice had increased numbers of lung ILC compared to C57BL/6J mice in both neonates and adults (Figure 4a-j). tSNE analysis revealed distinct clustering between the strains in both neonates (Figure 4k-l) and adults (Figure 4m-n). Neonatal BALB/c mice had reduced MFI of ST2, CD90.2, SCA-1 and KLRG1 and increased ICOS compared to C57BL/6J mice (Figure 4o-s). Adult BALB/c mice had lower expression of ST2 and SCA-1, increased levels of CD90.2 and ICOS, and similar levels of KLRG1 compared to C57BL/6J mice (Figure 4t-x).

**Figure 4:**
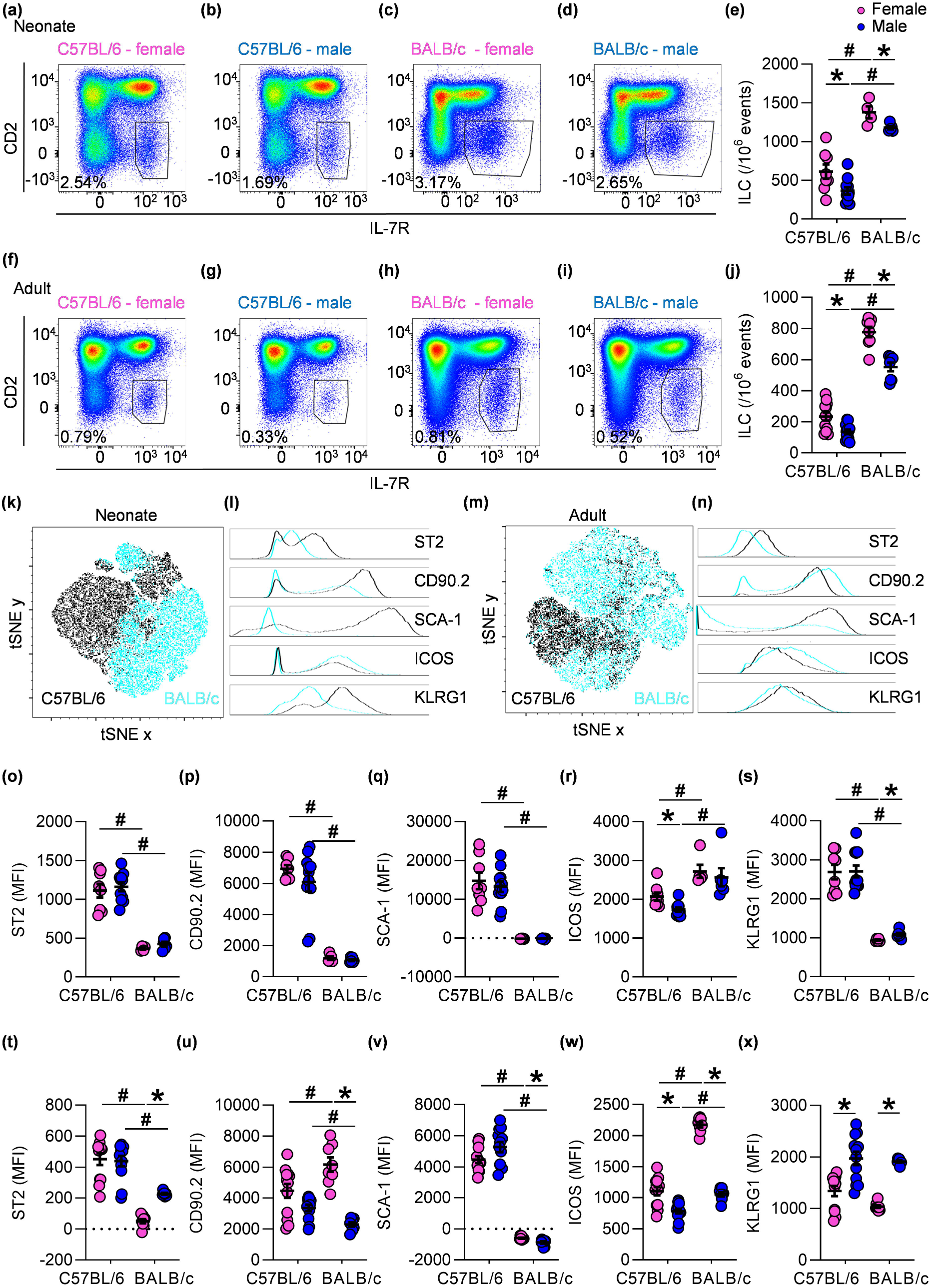
BALB/c mice have increased numbers of lung ILC and differential expression of ILC2-associated cell surface markers compared to C57BL/6J mice. **(a-d)** Representative pseudocolour plots of CD45^+^ lymphocytes shown as CD2^+^ *versus* IL-7R^+^ from **(a)** female C57BL/6J neonates (postnatal day 10), **(b)** male C57BL/6J neonates, **(c)** female BALB/c neonates and **(d)** male BALB/c neonates. **(e)** Quantification of **(a-d)**. **(f-i)** Representative pseudocolour plots of CD45^+^ lymphocytes shown as CD2^+^ *versus* IL-7R^+^ from **(f)** female C57BL/6J adult (7-9 weeks), **(g)** male C57BL/6J adult, **(h)** female BALB/c adult and **(i)** male BALB/c adult lungs. **(j)** Quantification of ILC in adult lungs. **(k)** Dimensionality reduction of neonatal ILC from BALB/c and C57BL/6J mice regarding ST2, CD90.2, SCA-1, ICOS and KLRG1 using t-Distributed Stochastic Neighbor Embedding (tSNE). tSNE x and tSNE y represent the two new parameters. **(l)** Histograms of ILC2-associated cell surface antigens. **(m)** tSNE of ILC from BALB/c and C57BL/6J adult mice. **(n)** Histograms from **n**. **(o-s)** Mean fluorescent intensity (MFI) of ILC cell surface markers in neonatal and **(t-x)** adult mice from data shown in **a**. Pink female, blue male mice. Data are presented as the mean ± the standard error of the mean (s.e.m). Data are representative of 2-4 independent experiments. Statistical significance was calculated using non-parametric unpaired Mann-Whitney t test. * P<0.05.

## Discussion

Pulmonary ILC2 are a heterogeneous lymphocyte population present in increased numbers during the first two weeks of life in the mouse lung^5–7,12^. Here we identify differences in pulmonary ILC numbers, ILC2-associated cell surface antigen expression, IL-5 and IL-13 expression and strain-dependency. Interestingly, we demonstrate that sex-dependent differences in pulmonary ILC occur in the neonatal period.

ILC progenitors, ILC2 and ILC3, are present in increased numbers during the first two weeks of postnatal lung development, with the increase in lung resident ILCs occurring concurrently with changes in lung stromal cells^5–7,10–12^. Lung alveolar fibroblasts provide a niche promoting the proliferation and maturation of ILC progenitors driven by insulin-like growth factor^10,11^. Adventitial stromal cells in the lung, expressing both IL-33 and thymic stromal lymphopoietin, have also been shown to support and regulate ILC2 proliferation^16^. Collectively, these data suggest that ILC activity is labile in the neonatal period of postnatal lung development. It remains unknown whether the interaction between neonatal lung ILC and stromal cells occurs in a bi-directional manner.

ILC2 are known secretors of the type 2 cytokines IL-5 and IL-13, and it has previously been shown that neonatal mice express more of these cytokines than adult mice^5,7,9,12,13,17^. This is further supported by our findings, which showed increased lung IL-5^+^ and IL-13^+^ lymphocytes in neonates compared to adults.

The current study compares cell numbers and cytokine expression, and the expression of ILC2-associated cell surface markers between sexes and ages. Whilst several studies have identified high plasticity in ILC^2,17–19^, with a recent report identifying two distinct ILC2 populations in both neonatal and adult lungs^17^, our study shows that markers associated with ILC2 activation, such as ST2 or ICOS, display varied expression during postnatal lung development. The neonatal period appears to be an important timeframe for expansion, activation and shaping of ILC2 signatures^12,13^. Lung ILC2 from the postnatal period are the main contributor to the adult lung ILC2 pool, and interestingly, ILC2 derived from the neonatal period are more responsive to stimulation with IL-33 in adulthood than ILC2 that were derived in adulthood^12,13^.

In adult mice, a sex-bias exists, with female mice having higher numbers of ILC^20–22^. Female mice have a specific KLRG1^−^ ILC2 population, which is not present in males^22^. These studies also showed phenotypic changes, with males expressing more KLRG1 and ST2 than females^20,22^. Castration of male mice increased ILC2 frequency, suggesting that androgen signalling negatively regulates ILC2, especially the KLRG1^−^ population^20–22^. While these studies did not observe differences in infant 3-week-old mice^21,22^, it should be noted that the current study found sex-specific differences much earlier in development at P10. In contrast to previous studies, our study found differences in ILC numbers and cell surface antigen expression as early as P10, with females having more ILC and higher ICOS expression than males. tSNE clustering of cell surface markers revealed more distinct clusters in adult compared to neonatal mice, indicating greater differences between sexes in adult mice.

Entwistle *et al.*, recently investigated lung ILC2 phenotypes in relation to strain, location and stimuli^23^. They could not detect significant changes in ILC2 numbers between BALB/c and C57BL/6J mice, and CD90.2 appeared to be the most stable marker between strains and stimuli^23^. In our current study, different ages, sexes and strains were compared at the ILC and not the ILC2 level, and increased CD90.2 expression was found in neonates compared to adults, and in female compared to male BALB/c mice, hence not making it a stable marker. Additionally, the present study detected more ILC in BALB/c compared to C57BL/6J mice in both neonatal and adult lungs.

The heterogeneity of pulmonary ILC makes it difficult to find reproducible cell surface antigens to identify a specific ILC subset. Our current study shows that lung ILC differ depending on mouse age, sex and strain, and highlights the need for caution regarding gating strategies and panel design. While there may not be a combination of cell markers that clearly identifies ILC or ILC2 in all situations, the field should aim to identify the most relevant and stable markers in specific targeted studies and be aware of the possibility that different cell surface antigen gating may yield variable results influenced by mouse age, sex and strain. The inclusion of reporters marking intracellular cytokines e.g. IL-5 and IL-13 or transcription factors such as Gata3 or RORα may aid the identification of specific subsets^14,15,24,25^.

This study shows increased numbers of lung ILC in neonatal compared to adult mice that warrants further exploration into the link between pulmonary ILC subsets and postnatal lung development. It adds new information to the field showing that sex can influence pulmonary ILC in the neonatal period, that IL-5 and IL-13 dual reporter mice are an effective tool for identifying ILC in the neonatal lung and that the strain of mouse has a clear impact on ILC number and cell surface antigen expression.

## Methods

### Mice

Pregnant female wild-type (WT; C57BL/6JAusb, BALB/cJAusb), 6-8-week-old female and male WT and IL-5/IL-13 dual reporter mice (*Il5*^venus/+^*Il13*^td-tomato/+^)^14,15^ were obtained from Australian Bioresources (Moss Vale, Australia). All mice were housed in individually ventilated cages under specific pathogen free physical containment 2 conditions. Mice were kept on a 12-hour day/night cycle with access to standard laboratory chow and water *ad libitum*. Before the start of experiments, adult mice had a 1-week acclimatisation period.

### Animal models

For the time-course study, neonates were euthanised at different time-points, with postnatal day (P) 0 being the date of birth. From P4 onwards, animal sex was determined. Adult mice (6-8 weeks) had one week of acclimatisation before euthanising for experiments.

### Flow cytometry

Lungs were collected from mice, ensuring no lymphatic tissue was attached, and single-cell suspensions were prepared according to the manufacturer’s instructions (Miltenyi Biotec GmbH, 2008). Cells were blocked with CD16/CD32 Fc block (BD Biosciences, 553141) for 30 minutes and stained with cell maker-specific antibodies (Supplementary Table 1) for 20-30 minutes. Staining and washing steps were performed with BSA stain buffer (554657, BD Biosciences). Samples were acquired on a BD FACSAria III (time-course figure 1) or BD LSR Fortessa flow cytometer (all other data) using FACSDiva software versions 8.01 and 9.01 (BD Biosciences). Unstained cells and single-stained compensation beads (BD Biosciences, 552843) were used for compensation of samples. Generally, 3-5×10^6^ events were acquired for each sample.

### Flow cytometric analysis and tSNE

Analysis of flow cytometry data was done using FlowJo versions 10.6 and 10.7 (BD Biosciences). ILC were gated as CD45^+^ single (FSC-A vs FSC-H) TCR^−^(TCRβ^−^TCRγδ^−^CD4^−^ CD8^−^) Lin^−^(CD11b^−^B220^−^Ly6a/e^−^NK1.1^−^CD3^−^Ter-119^−^) CD2^−^IL-7R^+^ cells. ILC were represented per 10^6^ total events acquired. For each tSNE clustering analysis, all compared samples were used in one workspace, and the compensation from one of the days applied to all samples. To not introduce bias, compensation .fcs files were chosen from a random day, and a new compensation matrix was adjusted and applied using FlowJo. Due to different expression intensities of markers used for gating and the need for identical gates for tSNE analysis, gates were adjusted in such a way that they covered the populations of interest in all of samples. Final ILC gates were concatenated into one .fcs file. The concatenated files were analysed with tSNE regarding parameters that were not used for the gating of ILC, and with the iterations 1,000, perplexity 30, using opt-SNE. Dual reporter mice were not used for tSNE analysis due to their endogenous cell fluorescence.

### Statistical analysis

All statistical analysis was done using GraphPad Prism 8.4.3. A maximum of one statistical outlier per experimental group was identified using the programme’s Grubbs test with alpha=0.05 and removed. Grubbs test was applied to every data set. For comparison of two groups, unpaired non-parametric Mann-Whitney test was used to compare ranks. Confidence interval was set at standard 95% or *P<0.05.

## Acknowledgements

MRS was supported by funding from Australian Research Council (DE170100226), National Health and Medical Research Council (NHMRC, APP1156898), Hunter Medical Research Institute, The University of Newcastle, Thoracic Society of Australian and New Zealand and Monash University. PMH is funded by Fellowships and grants from the NHMRC (1079187, 1175134) and by the University of Technology Sydney, University of Newcastle. GTB is funded by an NHMRC Fellowship (1135898). IL-5 reporter mice were kindly provided by Prof. Paul Foster, The University of Newcastle, Australia. IL-13 reporter mice were kindly provided by Prof. Andrew McKenzie MRC Laboratory of Molecular Biology, Cambridge, United Kingdom.

