## Supplemental methods for "Differences in pulmonary innate lymphoid cells are dependent on mouse age, sex and strain"

Short running title: Age, sex and strain impact lung ILC

Svenja Loering^1^, Guy J. M Cameron^1^, Nirmal P Bhatt^1^, Gabrielle T Belz^2, 3^, Paul S Foster^1^, Philip M Hansbro^1, 4, #^, Malcolm R Starkey^1, 5, #, *^

Author Affiliations

^1^Priority Research Centre’s GrowUpWell and Healthy Lungs, School of Biomedical Sciences and Pharmacy, Faculty of Health and Medicine, The University of Newcastle and Hunter Medical Research Institute, Newcastle, New South Wales, Australia

^2^The University of Queensland Diamantina Institute, Faculty of Medicine, The University of Queensland Translational Research Institute, Woolloongabba, Queensland, Australia

^3^Division of Molecular Immunology, Walter and Eliza Hall Institute of Medical Research, Parkville, Victoria, Australia

^4^Centre for Inflammation, Centenary Institute and University of Technology, School of Life Sciences, Faculty of Science, Sydney, New South Wales, Australia

^5^Department of Immunology and Pathology, Central Clinical School, Monash University, Melbourne, Victoria, Australia

^#^authors contributed equally

**Supplementary data**

**Supplementary Table 1: Antibodies used for flow cytometric identification of lung ILC**

| Antibody | Order number | Clone | Fluorophore | Assay dilution | Source |
| --- | --- | --- | --- | --- | --- |
| CD278 (ICOS) | 565886 | C398.4A | BV421 | 100 | BD Biosciences |
| TRC-β | 563221 | H57-597 | BV510 | 300 | BD Biosciences |
| TCR-γδ | 563218 | GL3 | BV510 | 300 | BD Biosciences |
| CD4 | 563106 | RM4-5 | BV510 | 300 | BD Biosciences |
| CD8a | 563068 | 53-6.7 | BV510 | 300 | BD Biosciences |
| KLRG1 | 740553 | 2F1 | BV650 | 100 | BD Biosciences |
| CD2 | 740655 | RM2-5 | BV711 | 300 | BD Biosciences |
| CD45 | 550994 | 30-F11 | PerCP-Cy5.5 | 200 | BD Biosciences |
| Ly-6A/E (Sca-1) | 565397 | D7 | BB515 | 50 | BD Biosciences |
| CD127 (IL-7R) | 562419 | SB/199 | PE-CF594 | 100 | BD Biosciences |
| CD25 | 552880 | PC61 | PE-Cy7 | 50 | BD Biosciences |
| IL-33R (ST2) | 17-9335-82 | RMST2-2 | APC | 50 | eBioscience |
| CD11b | 557960 | M1/70 | AF700 | 100 | BD Biosciences |
| Ly-6G/C (Gr-1) | 557979 | RB6-8C5 | AF700 | 100 | BD Biosciences |
| CD45R/B220 | 557957 | RA3-6B2 | AF700 | 100 | BD Biosciences |
| Ter-119/Erythroid cells | 560508 | TER-119 | AF700 | 100 | BD Biosciences |
| CD3 molecular complex | 561388 | 17A2 | AF700 | 100 | BD Biosciences |
| NK-1.1 | 560515 | PK136 | AF700 | 100 | BD Biosciences |
| CD90.2 | 561641 | 53-2.1 | APC-Cy7 | 200 | BD Biosciences |
| *Il5*^venus^ |  |  | BB515 |  |  |
| *Il13*^td-tomato^ |  |  | PE |  |  |
